# Protein aggregation in plant mitochondria inhibits translation and induces an NAC017-dependent ethylene-associated unfolded protein response

**DOI:** 10.1101/2023.01.11.523570

**Authors:** Ce Song, Yuanyuan Li, Yuqi Hou, Mengmeng Yang, Tiantian Li, Yinyin Liu, Chang Xu, Jinjian Liu, A. Harvey Millar, Ningning Wang, Lei Li

**Affiliations:** Frontiers Science Center for Cell Responses, Department of Plant Biology and Ecology, College of Life Sciences, Nankai University, 300071 Tianjin, China; Key Laboratory of Radiopharmacokinetics for Innovative Drugs, Chinese Academy of Medical Sciences, and Institute of Radiation Medicine, Chinese Academy of Medical Sciences & Peking Union Medical College 300192 Tianjin, China; ARC Centre of Excellence in Plant Energy Biology, School of Molecular Science, The University of Western Australia, 6009 Crawley, WA, Australia

**Keywords:** mitochondrion, UPR, PPR, ethylene

## Abstract

Loss of Lon1 in plant mitochondria led to stunted plant growth and accumulation of nuclear-encoded mitochondrial proteins, including Lon1 substrates, while mitochondrial-encoded proteins typically decreased in abundance. Lon1 mutants contained protein aggregates in the mitochondria matrix which were enriched in PPR-containing proteins and ribosomal subunits of the translation apparatus and were slowed in mitochondrial RNA splicing, editing and general translation rate. Transcriptome analysis showed multiple organellar unfolded protein responses involving ethylene biosynthesis were induced by either Lon1 loss, mitochondrial ribosomal protein loss, translation or respiratory inhibition and most were regulated by the mitochondrial retrograde signaling pathway dependent on the transcription factor NAC017. The short hypocotyl in *lon1* mutants during skotomorphogenesis was partially rescued by ethylene inhibitors and mutants showed higher ethylene production rates than wildtype. Together this provides multiple steps in the link between loss of Lon1 and its whole plant phenotype.

**Single Sentence Summary:** Lon1 knockout inhibits mitochondrial-encoded gene translation and induces retrograde signaling involving unfolded protein responses.

## Introduction

Mitochondria generate cellular ATP and are involved in the synthesis of amino acids, vitamins and reactive oxygen species throughout the plant life cycle. Coordination of nuclear-encoded protein import and *in organello* protein synthesis from the small number of mitochondrial-encoded proteins is needed to ensure subunit stoichiometry in electron transport chain (ETC) protein complexes and 70S ribosomes which contain proteins encoded in both genomes. In mitochondria, a network of protein quality control systems involving proteases and chaperones work together for the correct folding, assembly and degradation of mitochondrial proteins to enable this to occur (van Wijk, 2015). Amongst them is Lon1, a dual function enzyme with both protease and chaperone domains that is located in the mitochondrial matrix in most eukaryotes, including plants (Rep et al., 1996; Dikoglu et al., 2015; Li et al., 2017b).

Lon (originally named La) was first characterised as a heat shock protein in *E*.*coli* almost half a century ago (Chin et al., 1*98*8). Later studies in eukaryotes found a Lon1 ortholog was located in mitochondria and highly conserved in yeast, plants and animals. Lon1 sequences and activity are highly conserved, for example the Arabidopsis Lon1 ortholog can complement the yeast Lon1 (PIM1) mutant phenotype (Rigas et al., 2009). Lon1 loss of function mutations cause obvious mitochondrial malfunction and morphology changes and lethality or severe growth and development retardation in both yeast and humans (Suzuki et al., 1994; Strauss et al., 2015). The yeast Lon1 mutant cannot survive in non-fermentation media, indicating it is required for mitochondrial function. Due to null mutations causing lethality, knock down mutants of animal Lon1 (LonP1) are often utilised to study its role and define its substrates (Pareek et al., 2018; Zhao et al., 2022). In contrast, knockout mutants of Lon1 in Arabidopsis are viable, but show growth retardation under normal conditions and low germination rate following heat treatments (Rigas et al., 2009).

In spite of these phenotypic differences, common morphological and mechanical mitochondrial changes are found in *lon1* mutants across organisms. Electron dense bodies which have been proposed to be protein aggregates have been observed under transmission electron microscopy in *lon1* mitochondria in yeast, Arabidopsis and humans (Suzuki et al., 1994; Rigas et al., 2009; Strauss et al., 2015). Decrease amounts of assembled ETC protein complexes and accumulation of TCA cycle enzymes have also been reported in all three organisms. Decrease in abundance of mitochondrial-encoded proteins due to Lon1 protease knockdown has been reported in Drosophila, murine and human cell lines (Pareek and Pallanck, 2018; Pareek et al., 2018; Zurita Rendon and Shoubridge, 2018; Zhao et al., 2022). Ribosome biogenesis inefficiency or inability of ribosome to interact with RNA have also been discovered in knock down mutants in Drosophila and a human cell line (Pareek et al., 2018; Zurita Rendon and Shoubridge, 2018). In contrast to the bacterial and human Lon1 oligo hexamers structure (Shin et al., 2021), plant Lon1 is predicted to form a heptamer due to its extended N terminus, similar to yeast (Rigas et al., 2014). A well-known Lon1 proteolytic substrate mitochondrial transcription factor A (TFAM) claimed to be important for its role in animals (Matsushima et al., 2010) is also missing in plants (Kim et al., 2021).

To investigate the role of Lon1 in plants and how they survive in its absence, the mitochondrial proteomes of two *lon1* mutant alleles (*lon1-1* and *lon1-2*) have previously been investigated by 2-D fluorescence difference gel electrophoresis (DIGE) showing a common accumulation of chaperones in addition to ETC protein complexes reduction (Solheim et al., 2012). Later protein turnover analysis found Lon1 loss cause large scale changes in mitochondrial protein turnover (Li et al., 2017b), indicating a very wide impact by Lon1 disruption on mitochondrial protein homeostasis.

In this study, we have sought to link protein aggregation, changes in mitochondrial function and transcriptional responses in *lon1* mutant alleles in an alternative way to explain whole plant phenotypes. To this end, proteins in mitochondrial aggregates were enriched, identified and quantified alongside the total mitochondrial proteome by peptide mass spectrometry. We found pentatricopeptide repeat (PPR) proteins involved in mitochondrial mRNA splicing and editing were highly enriched in protein aggregates. We then showed that *lon1* mitochondria were functionally limited in RNA splicing, editing and translation. This limitation correlated with specific reduction in the abundance of mitochondrial-encoded proteins compared to nuclear-encoded mitochondrial proteins. We also found *lon1* mutants induce an unfolded protein response similar to that observed following mitochondrial ribosomal protein loss, translation loss or respiratory inhibition. The genes involved include ethylene signaling factors and are shown to be regulated by the transcription factor NAC017 that controls a key pathway in mitochondrial retrograde signaling in plants. We show elevated ethylene signaling in *lon1* mutants to be at least partially responsible for the *lon1* phenotype, evidenced by ethylene measurements and partial rescue of the short hypocotyl in *lon1* mutants during skotomorphogenesis by ethylene inhibitors.

## Results

### The abundance of mitochondrial-encoded proteins is depleted in *lon1* mutants

Two Lon1 loss of function lines (EMS induced point mutation line *lon1-1* and T-DNA insertion line *lon1-2*) and *Col-0* control plants were grown on half-strength MS plate and/or grown in soil for phenotyping (**Fig 1A**). Short root, small rosette and growth retardation in *lon1* mutants were observed in the seedling and vegetative growth stages, as reported previously (Rigas et al., 2009; Solheim et al., 2012). Gel-based (DIGE and SDS-PAGE) and iTRAQ-based proteomics analysis were previously performed to investigate protein abundance or protein turnover changes in *lon1* mutants, but were limited in their coverage of mitochondrial-encoded proteins (Solheim et al., 2012; Li et al., 2017b). An in-depth label-free quantification (LFQ) was therefore performed here using mitochondria extracted from Arabidopsis *Col-0* and two *lon1* mutant lines. In total, the abundance of 893 and 8*98* proteins could be measured in *lon1-1* and *lon1-2* and compared to *Col-0*, respectively (**Fig 1B, DataS1**). Over half the proteins showed statistically significant accumulation in two *lon1* mutants, while less than 15% of proteins showed statistically significant reduction (Student’s T-test, P<0.05). To explore changes in the stoichiometry between nuclear and mitochondrial genome encoded proteins, a two-sample Kolmogorov-Smirnov test was used to compare proteins in *lon1* mutants and *Col-0* (**Fig 1C**). This showed that the majority of the thirteen mitochondrial-encoded proteins that were identified in samples decreased in abundance in *lon1* mutants (**Fig 1D**). Complex I subunits (NAD1, 2, 4, 5, 6, 7, 9), cytochrome c maturation associated protein CCB452, Complex IV subunit COB and Complex V (i.e. ATP synthase) subunit ATPb all decreased in abundance. In contrast, ribosomal subunits (RPS3, 4) and an ATP synthase subunit ATP6 increased in abundance. These common changes, mostly of which were decreases in protein abundance, indicate a general decrease in protein homeostasis for mitochondrial-encoded genes.

**Fig 1.**
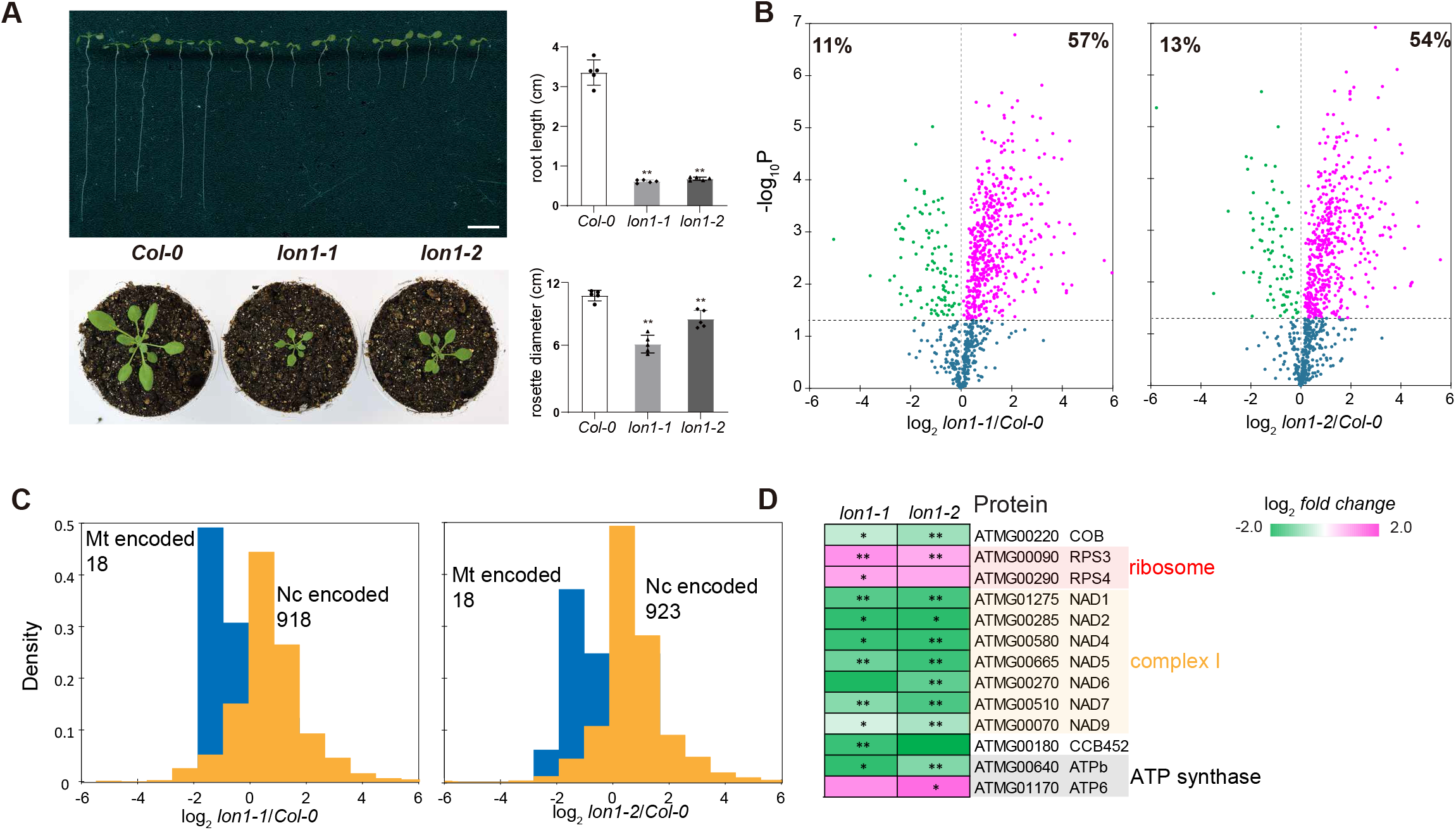
Arabidopsis Lon1 loss of function leads to growth retardation and decreases in abundance of mitochondrial genome encoded proteins. Root length and rosette diameter of 10-day-old plate grown Arabidopsis seedlings and 21-day-old soil grown plants were measured and compared between *Col-0* and two *lon1* lines **(A)**. Protein abundance of extracted mitochondria from *Col-0* and *lon1* mutant lines were measured by a label free quantification (LFQ) mass spectrometry strategy. Log-transformed fold changes in abundance of 893 and 8*98* proteins between *lon1-1/lon1-2* and *Col-0* and shown as volcano plots. Percentage of statistically significant changes were shown **(B)**. Nuclear (Nc) and mitochondrial (Mt) encoded proteins were plotted as separate histograms. Two-sample Kolmogorov-Smirnov test was performed to determine the P values between two distributions (P<0.01) **(C)**. Log-transformed fold changes in abundance of thirteen proteins with statistically significant changes were shown as heatmaps **(D)**. A Student’s T-test was performed to determine statistical significance (**A** and **D**\*\* means P<0.01, * means P<0.05).

### Mitochondrial aggregates are enriched in metabolic enzymes and proteins involved in translation and protein homeostasis

Transmission electron microscopy has observed protein aggregates in the mitochondrial matrix of *lon1* mutants in yeast, Arabidopsis and mammals (Suzuki et al., 1994; Rigas et al., 2009; Strauss et al., 2015). To study their contents in Arabidopsis, they were enriched based on their low solubility by a nonionic detergent Triton X-100 (Wilkening et al., 2018; Maziak et al., 2021; Pollecker et al., 2021). Proteins from total mitochondria, and detergent-solubilized supernatant and protein aggregates were separated by SDS-PAGE and stained for visualisation (**FigS1**). The pattern of protein bands in the precipitate fraction were distinct in *lon1* lines compared to *Col-0*. Some changes in specific band patterns were also observed in total mitochondrial fractions but not in the detergent-solubilised supernatant fractions. The similarity in protein banding patterns between *lon1* mutant lines and among biological replicates suggested the same proteins tend to form aggregates in the mutants. Using a label free quantification strategy, we measured the abundance of 459 proteins and compared them between the precipitate fractions from *Col-0* and *lon1* mitochondria (**DataS2**). Compared with *Col-0*, 169 and 173 proteins were significantly different in abundance in the precipitant fraction of *lon1-1* and *lon1-2*. To compensate for proteins which changed in total mitochondrial abundance in mutants, we normalized abundance in the precipitant fraction to the total mitochondrial protein abundance to calculated an aggregation factor (AGF, pellet-FC/total-FC) for a specific protein. Fifty-seven proteins had a mean AGF value greater than 1.5 in both *lon1* mutant lines (**Fig 2, DataS3**). Two groups of protein stood out in this list, metabolic enzymes of the TCA cycle and amino acid metabolism, and translation and protein stability machinery proteins PPR/transcription, ribosomal subunits and proteases. The presence of 16 subunits of TCA cycle enzymes and 18 subunits of amino acid metabolism enzymes is consistent with previous evidence that a range of TCA enzymes declined in activity but increased in total mitochondrial abundance in *lon1* lines (Rigas et al., 2009; Solheim et al., 2012; Li et al., 2017b). Aggregated copies of these enzymes are likely to be inactive (Bender et al., 2011; Li et al., 2017b). But the reason for the group of PPR and ribosomal proteins participating in mitochondrial-encoded gene transcription and translation in protein aggregations (**Fig 2A**) required further investigation.

**Fig 2.**
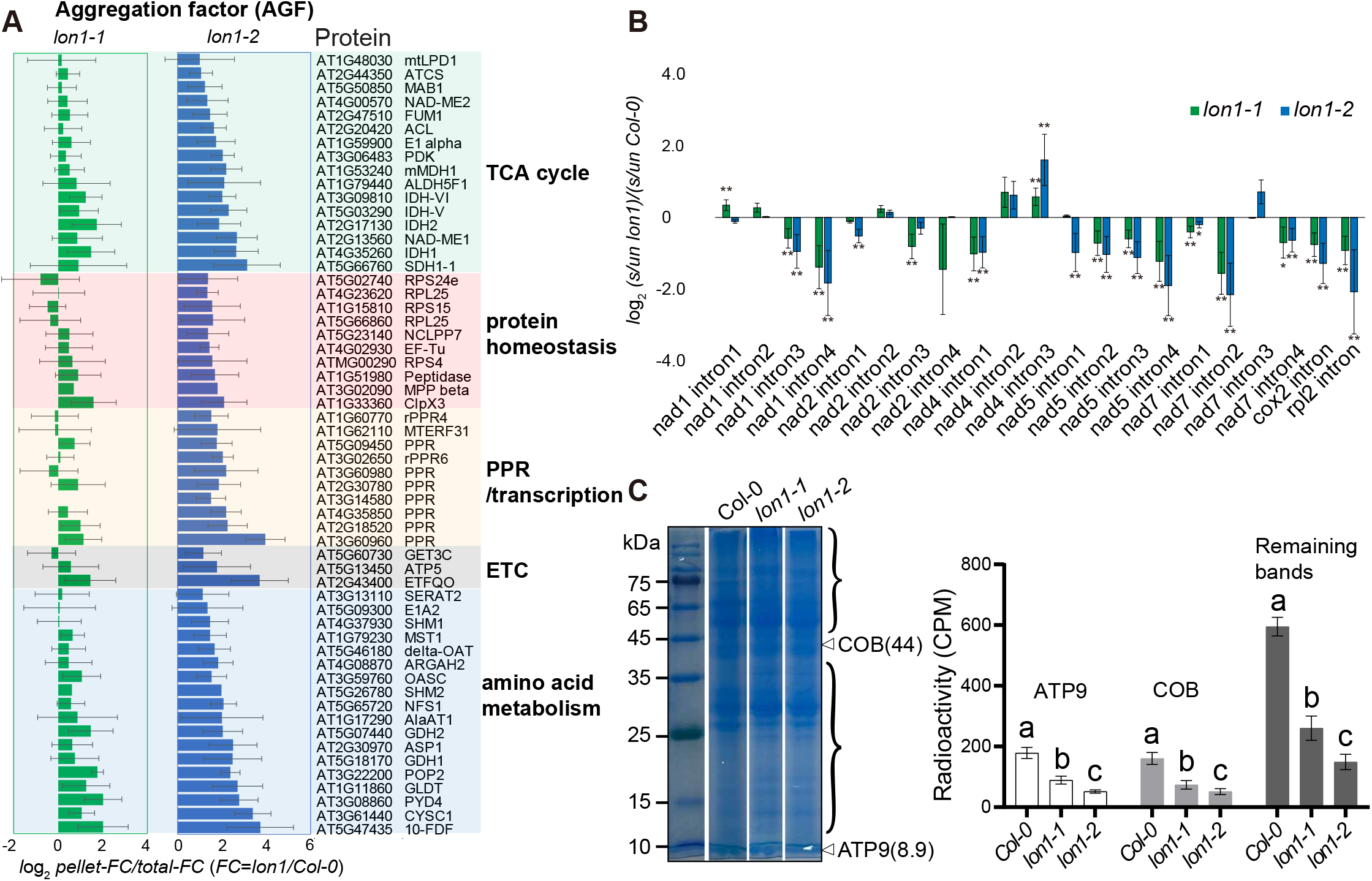
Protein aggregates containing PPR and ribosomal proteins led to deficient mitochondrial splicing and translation in *lon1* mutants. The top 5 major functional GO categories which are enriched in *lon1* protein aggregates. Fifty-seven proteins with statistically significant accumulation in at least one *lon1* mutant lines based on average aggregation factors over 1.5 fold (**A**). Spliced (s) and unspliced (un) transcripts for mitochondrial-encoded proteins were quantified by qRT-PCR and plotted as log-transformed fold changes of *lon1* mutants compared to *Col-0* (**B**). Mitochondria used for in organello protein synthesis assays were fractionated by 12% SDS-PAGE (**C**). Mitochondrial-encoded ATP synthase subunit ATP9 and Complex III COB bands (triangle arrows) were cut separately for radioactivity assays. Radioactivity of ATP9, COB and remaining protein bands containing [^35^S]-Methionine were measured by liquid scintillation counting as counts per minute (CPM). One-way ANOVA was performed for statistical analysis between *Col-0* and *lon1* mutant lines for the same bands.

### *lon1* mutants show slow or inefficient RNA splicing, editing and translation inside mitochondria

To determine how aggregation of PPRs and ribosomal proteins might affect their function, we performed RNA splicing, editing and in organelle translation assays. Spliced and unspliced transcripts were quantified by qRT-PCR using promoters reported previously (Zhao et al., 2020). The splicing efficiency of seven mitochondrial-encoded genes (*nad1, 2, 4, 5* and *7, cox2* and *rbl2*) in Arabidopsis seedlings were determined and compared between *Col-0* and *lon1* lines. Most tested genes decreased in splicing efficiency with statistical significance, with a few exceptions, namely *nad1* intron 1 and *nad4* intron 3 (**Fig 2B**). On the other hand, the expression level of fifteen mitochondrial-encoded genes rose roughly two fold in *lon1* mutant lines, suggesting either stimulated mitochondrial transcription or a potential increase of mitochondrial genome number (**FigS3A**). To evaluate RNA editing in mitochondria, seventy sites of known C-U RNA editing (Guillaumot et al., 2017) were determined and measured by cDNA sequencing (**FigS3B, DataS4**). Compared with *Col-0*, fifteen known C-U RNA editing sites (Complex I-*nad1, 2, 5 and 7*; Complex V-*atp1, atp4, atp6-1, atp6-2 and atp9*) decreased in *lon1* mutants. Furthermore, we performed radioactive [^35^S]-Met in organelle protein synthesis assay to investigate translation efficiency in *lon1* lines (**Fig 2C**). Radioactivity was measured by liquid scintillation counting as counts per minute (CPM). The two most abundant mitochondrial-encoded proteins, ATP9 and COB, were sliced separately from gels, while all remaining proteins were pooled together for quantification of the radioactivity assay. Radioactivity of all three fractions showed a general decrease in *lon1* lines suggesting deficient translation without functional Lon1.

### Unfolded protein response, ethylene biosynthesis and signaling genes are upregulated in *lon1* mutants

Inhibition with mitochondrial translation inhibitor doxycycline (DOX) or deficient mitochondrial translation due to a ribosomal subunit mutation had been found to induce mitochondrial unfolded protein response (UPR^mt^) (Wang and Auwerx, 2017). Mitochondrial protein aggregations in *lon1* mutants might therefore induce UPR^mt^ as a means to recover protein folding and functions. As a preliminary test of this hypothesis we found that DOX treatment could reproduce the *lon1*-like short root and growth retardation phenotype in *Col-0* (**FigS5**). To more formally assess the hypothesis, RNA sequencing was performed to investigate the transcriptome response in one of the *lon1* mutants; *lon1-2*. Transcripts of 27,445 genes were quantified in *lon1-2* and *Col-0* (**DataS5**). GO enrichment was performed using 4662 upregulated genes (FC>2, P<0.05) as target and all encoding genes as background (**FigS4A-B**). Besides the apparent enrichment of genes encoding mitochondrial localised proteins, UPR associated genes encoding ‘de novo’ protein folding and unfolded protein binding proteins are highly enriched in *lon1-2*. We further investigated reported genes involving in mitochondrial UPR (UPR^mt^) (Wang and Auwerx, 2017; Tran and Van Aken, 2020), chloroplast UPR (UPR^cp^) (Dogra et al., 2019) and endoplasmic reticulum UPR (UPR^er^) (Kim et al., 2018) in *lon1-2*. To dissect the pattern of transcription responses in *lon1* mutant lines, we compare all significant changes in *lon1-2* to published transcriptome data on mitochondrial translation deficient responses (doxycycline treatment and *mrpl1*), enhanced mitochondrial retrograde signaling (antimycin A treatment), and overexpression and loss-of-function of the major retrograde signaling transcription factor ANAC017 (**Fig 3, DataS6**). UPR genes show general upregulation which indicate cellular UPRs in *lon1-2*. UPR^mt^ genes are upregulated in *lon1-2* which are similar to deficient mitochondrial translation, retrograde signaling antimycin A and *anac017* overexpression. Expression of UPR^mt^ target genes (*HSP70*-*1, HSP60*-*2, SLP*-*1, NAC017, AOX1a*) was further validated by quantitative real-time PCR in two *lon1* lines (**FigS6**). UPR^mt^ genes upregulation was largely inhibited in an *anac017* point mutation line *rao2*.*1* which suggest ANAC017 plays a dominant role in retrograde signaling involving UPR^mt^, including those induced in *lon1* (**Fig 3**). UPR^cp^ candidate genes were also specifically stimulated in *lon1-2* mutant with only a proportion of chaperones also upregulated in transcript datasets for *anac017* overexpression and *anac017* point mutation lines. Unlike UPR^mt^ and UPR^cp^ target genes, less than half of UPR^er^ target genes were upregulated in *lon1-2* (**DataS6**). Only a few UPR^er^ target genes involving *nac103, ERF105, STZ* (salt tolerance zinc finger) and *T6J4*.*21* were also upregulated under antimycin A treatment and the *anac017* overexpression line.

**Fig 3.**
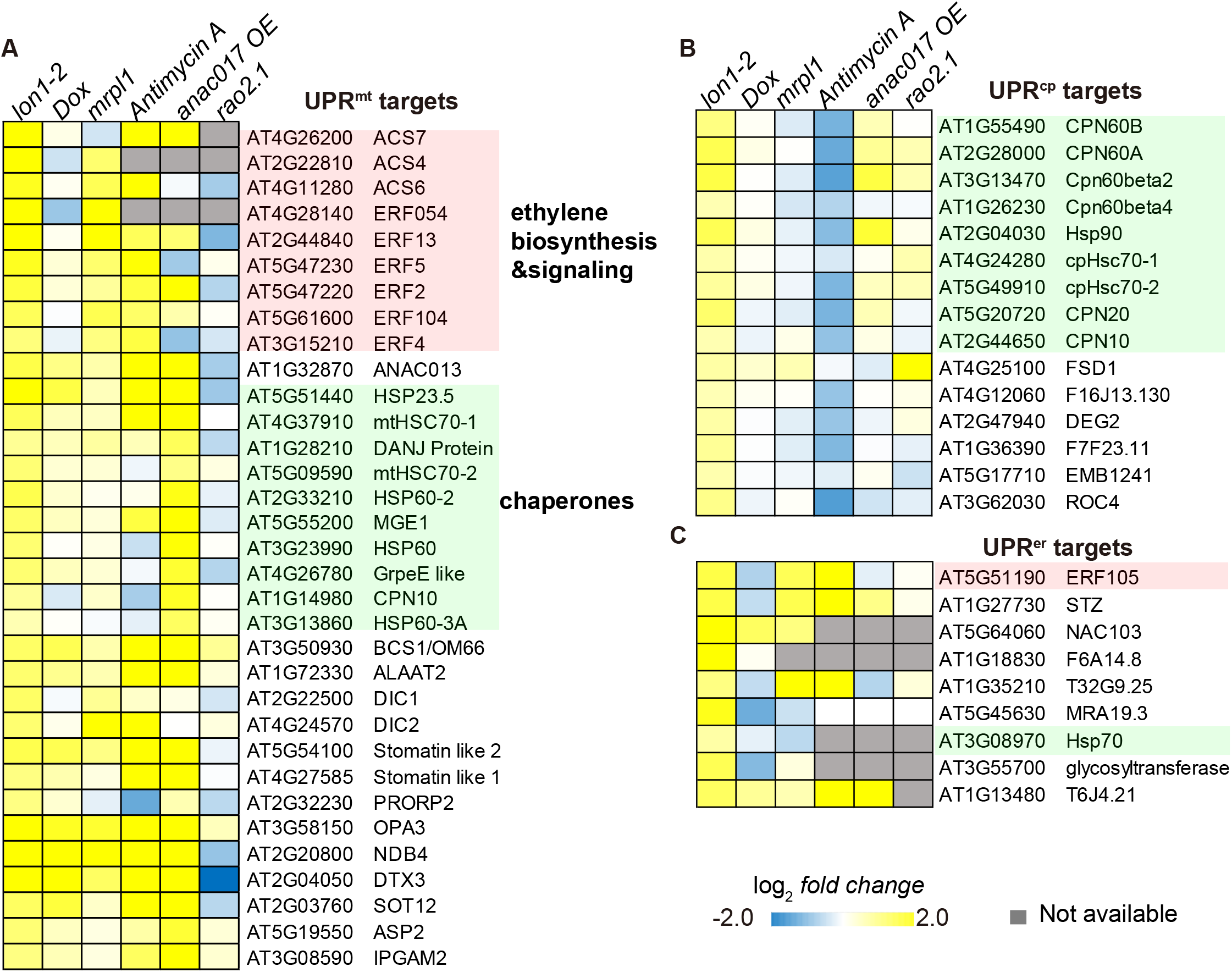
Arabidopsis Lon1 disruption induces unfolded protein responses and ethylene biogenesis. Transcripts for unfolded protein responses (UPR) involving UPR^mt^ (Wang and Auwerx, 2017; Tran and Van Aken, 2020), UPR^cp^ (Dogra et al., 2019) and UPR^er^ (Kim et al., 2018) target genes were induced in *lon1-2* (**A-C**). Transcripts with statistical significant induction (FC>1.5, P<0.05) in *lon1-2* was shown and compared with doxycycline (Dox) treatment, a *mrpl1* mutant (Wang and Auwerx, 2017), antimycin A treatment (He et al., 2022), *anac017* overexpression and the *anac017* point mutation line *rao2*.*1* (Meng et al., 2019).

### Etiolated *lon1* seedlings show an ethylene overactive phenotype that can be attenuated by ethylene inhibitors

As known target genes in UPR, ethylene synthesis and signaling genes were up-regulated in *lon1*. Key ethylene biogenesis enzymes *ACS4, ACS6* and *ACS7*, ethylene response factors *ERF2, ERF4, ERF5, ERF13, ERF54, ERF103* and *ERF104* were significantly up-regulated (**Fig 3A**). This guided us to consider if there was a role of ethylene biogenesis and signaling in establishing retrograde signaling in *lon1* mutants and explaining their phenotypes. We tested this by using measurements of the ethylene triple response; shortening and thickening of hypocotyls and roots and exaggeration of the curvature of apical hooks. It is known that ethylene acts as an inhibitor of hypocotyl elongation in darkness (Yu and Huang, 2017). Length of hypocotyl was measured in five-day-old etiolated Arabidopsis *Col-0* and *lon1* mutant lines. An ethylene overproduction mutant *eto1-12* and an ethylene-insensitive mutant *ers2-1* were used as positive and negative controls. Noticeably, hypocotyl elongation of *lon1-1, lon1-2*, and *eto1-12* was significantly inhibited under dark conditions compared to *Col-0* and *ers2-1* (**Fig 4A**). We further applied AgNO_3_ (ethylene receptor inhibitor) and AVG (ACC synthase inhibitor) to stop the ethylene signaling and production. Similar to *eto1-12*, hypocotyl elongation recovery was found in *lon1-1, lon1*-2 after AgNO_3_ and AVG treatment (**Fig 4B**). Apical hook curvature was measured and compared before and after AgNO_3_ and AVG treatments. Only *eto1-2* plants showed a larger apical hook curvature but not in other tested lines. AgNO_3_ and AVG treatments led to cotyledon opening with smaller apical hook curvature and accelerated the skotomorphogenesis to photomorphogenesis transition in all tested Arabidopsis lines. Hypocotyl diameter in *lon1* etiolated seedlings was thinner (**FigS7**). *Cell*ulose and lignin staining of hypocotyl transverse area showed lignin biogenesis was delayed in the root stele in *lon1* mutants. AgNO_3_ and AVG treatments diminished the observed difference between *Col-0* and *lon1* mutant lines, but through inhibiting the hypocotyl diameter and lignin biogenesis in *Col-0*, possibly independent of their ethylene inhibition effects.

**Fig 4.**
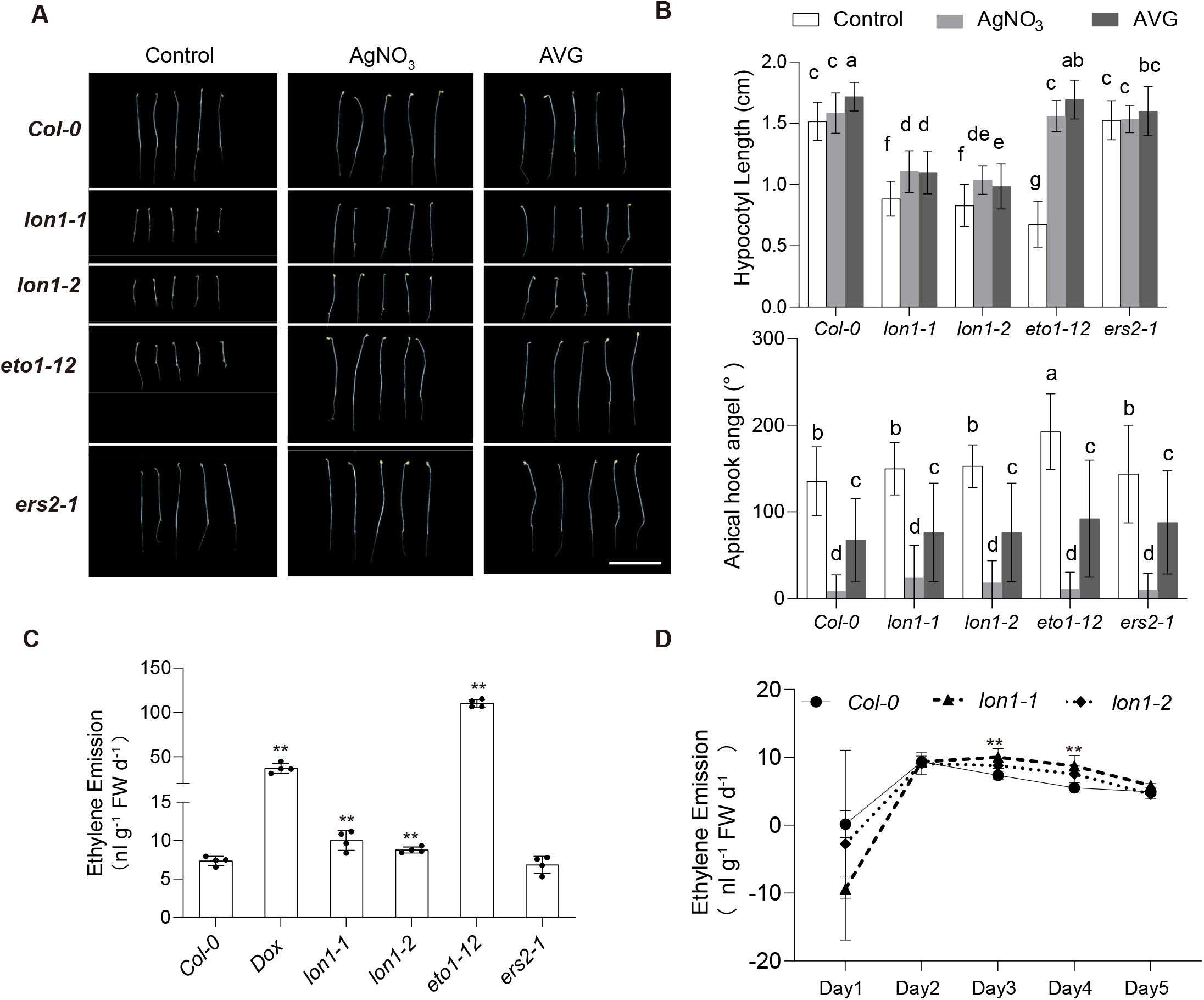
Lon1 disruption leads to short hypocotyl through more ethylene production in etiolated Arabidopsis seedlings. Five-day-old etiolated seedlings of *Col-0, lon1* mutants, Dox treatment and mutant lines with high (*eto1-12*) or no change in ethylene (*ers2-1*) (**A**). Hypocotyl length and apical hook angle of five-day-old plants under control conditions and after ethylene inhibitor AgNO_3_ and AVG addition are shown (**B**). One-way ANOVA was performed for groups shown. Ethylene emission from three-day-old etiolated seedlings was measured by gas chromatography and normalized to fresh weight (**C**). An ethylene emission curve over five days after sowing was shown for *Col-0* and *lon1* mutant lines (**D**). Error bars show standard deviations in four biological replicates. Statistical significance was determined by a Student’s T-test (**C** and **D**\*\* means P<0.01, * means P<0.05).

Due to an incomplete ethylene triple response to ethylene overproduction in *lon1*, we proceeded to measure the ethylene emission from *Col-0* and *lon1* mutant lines, using DOX treatment and *eto1-12* as positive control and *ers2-1* as negative control (**Fig 4C-D**). Ethylene emission from *lon1* mutants was significantly higher than in *Col-0*, while DOX-treated *Col-0* and *eto1-12* produced even more ethylene. Ethylene emission can change with early seedling development. We therefore measured the ethylene emission from *Col-0* and *lon1* mutant seedlings day one after sowing and continued for another five days. It was found that ethylene emission from *Col-0* and *lon1* mutant lines peaked at day two, but *lon1* lines emitted more ethylene for two days and then decreased to the same level as *Col-0* in darkness (**Fig 4D**).

## Discussion

### Lon1 controls abundance of mitochondrial-encoded protein through PPR and translation apparatus

Less cristae and changes in protein aggregation mitochondrial morphology were observed by transmission electron microscopy (TEM) in knockout or dysfunctional Lon1 in yeast, Arabidopsis and human cells (Suzuki et al., 1994; Rigas et al., 2009; Dikoglu et al., 2015). In this study, we isolated Arabidopsis mitochondria and enriched the mitochondrial protein aggregates for an integrated proteomics analysis (**FigS1**). Abundance of more than half the identified proteins altered in Lon1 knockout lines, while mitochondrial-encoded proteins showed a general decrease in protein abundance with few exceptions (**Fig 1B-D**). As a semi-autonomous organelle in the plant cell, the stoichiometry between nuclear and mitochondrial-encoded proteins is an important factor for mitochondrial protein homeostasis. Reduction in mitochondrial-encoded proteins, which are mostly protein subunits in electron transport chain (ETC) protein complexes (e.g. Complex I), can lead to inefficient protein complex assembly. Mitochondrial-encoded subunits of Complex I showed a general decrease in abundance. This provides a new explanation for the decreased assembled Complex I but accumulation of nuclear-encoded protein assembly intermediates in *lon1* lines that have been reported previously (Solheim et al., 2012; Li et al., 2017b).

Protein aggregation in *lon1* can inhibit metabolic enzyme activities (Rigas et al., 2009; Li et al., 2017b), but it has not been clear if other proteins found in aggregates such as those involved in protein homeostasis, indicated a similar effect on function (**Fig 2A, FigS2**). Namely proteins such as ribosomal subunits, translation regulators, proteases and PPR proteins. Ribosomal PPRs (rPPRs) are plant specific components of functional mitochondrial ribosomes (Waltz et al., 2019). As generic but essential plant translation factors, rPPR null mutations are either lethal or cause growth retardation. PPR-containing proteins also take part in RNA editing, splicing and translation. While nuclear transcription was upregulated in *lon1* lines, inefficient splicing of mitochondrial-encoded transcriptional products was evident from lower ratios of spliced to unspliced RNA (**Fig 2B**). Mild decreases in C-U sites were also observed in *lon1* lines (**FigS3**). Inefficient RNA splicing and editing can both contribute to less active transcripts for translation. Slower translation was also evident from an *in-organelle* translation assay in *lon1* mutants (**Fig 2C-D**). These three pieces of evidence together reveal that decreased mitochondrial-encoded protein abundance was consistent with post-transcriptional control involving aggregated and potential dysfunctional PPRs, translation factors and ribosomal proteins in Arabidopsis *lon1* lines (**Fig 5**). The wider list of newly identified proteins enriched in protein aggregates also provide an important resource to further investigate Lon1 functions and transcriptional regulation in plant mitochondria.

**Fig 5.**
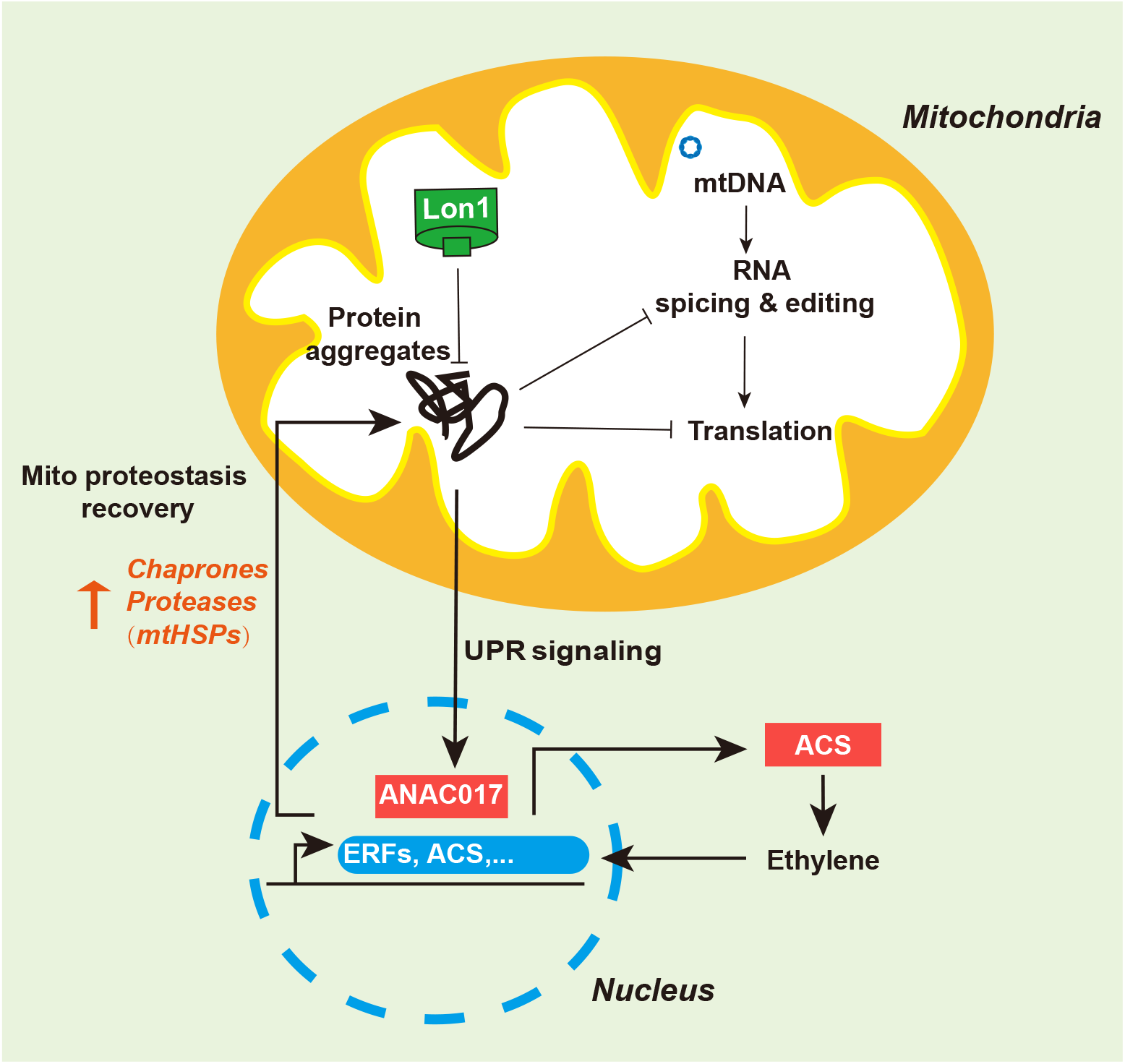
Lon1 prevents protein aggregations which inhibit translation and stimulate UPRs to rebalance mitochondrial proteostasis. Lon1 prevents protein aggregations which are enriched in PPRs, ribosomal subunits and metabolic enzymes. Aggregated PPRs are inefficient in RNA splicing and editing and lower in translation rate of mitochondrial-encoded mRNA. Reduction in mitochondrial-encoded protein synthesis contribute to less assembled electron transport chain complexes. Protein aggregation in *lon1* mutants induce ANAC017-dependent UPR signaling associated with ethylene overproduction. Ethylene stimulates the ERF transcription factors which activate UPR involving mitochondrial localised chaperones and proteases to rebalance proteostasis.

Decrease in abundance of mitochondrial-encoded proteins following Lon1 protease knockdown have also been reported in Drosophila, murine and human cell lines (Pareek and Pallanck, 2018; Pareek et al., 2018; Zurita Rendon and Shoubridge, 2018; Zhao et al., 2022). Ribosome biogenesis inefficiency or inability of ribosomes to interact with RNA was first discovered in Drosophila and a human cell line (Pareek et al., 2018; Zurita Rendon and Shoubridge, 2018). Here, the translation apparatus involving ribosomal subunits, rPPR and elongation factor (EF-Tu) was found to be enriched in mitochondrial aggregates which can inhibit active translation rates (**Fig 2A**). Moreover, deficient RNA splicing and editing can contribute to slower translation in the absence of Lon1 (**Fig 2B, Fig 5** and **FigS3**). In our view, this provides a more integrated explanation for the decrease in abundance of mitochondrial-encoded proteins in Arabidopsis and possibly also in other organisms as outlined above. PPR containing proteins were previously found to degrade slowly, but components of the translation apparatus including ribosomal subunits and rPPR6 were found to degrade more rapidly in *lon1* mutant (Li et al., 2017a; Li et al., 2017b). This suggests Lon1 may play a different role in maintenance of PPR-containing proteins and ribosomes which deserves investigation in future studies. In contrast to the majority of mitochondrial-encoded proteins, we found RPS3, RPS4 and ATP6 were more abundant in *lon1* lines (**Fig 1D**). This suggests a positive, selective control exists to accumulate these proteins in *lon1* lines.

### *Cell*ular unfolded protein responses occur in *lon1* by canonical and noncanonical signaling

The large-scale alteration in the mitochondrial proteome in *lon1* lines suggests a transcriptome response to rebalance mitochondrial protein homeostasis in the absence of protease. RNA sequencing analysis found dramatic transcriptional changes in a *lon1* mutant rich in mitochondrial-localised proteins (**FigS4, DataS7**). A correlation between mitochondrial protein abundance and corresponding transcript abundance showed an intermediate correlation (R=0.23), but over 70% of gene expression was positively correlated with protein abundance. This suggests that gene expression contributes to a large proportion of the protein abundance changes observed in *lon1* mitochondria (**Fig 1**). The group of genes with upregulated expression was enriched with protein folding, refolding or unfolded protein binding biological processes and molecular function categories, suggesting an upregulation of UPR. In fact, genes for cellular UPRs including UPR^mt^, UPR^cp^ and UPR^er^ were all upregulated in *lon1* (**Fig 3**).

UPR^mt^ in *lon1* was found to be induced by the canonical mitochondrial retrograde signaling pathway, regulated by ANAC017. This was support by the fact that the same list of target genes was stimulated by mitochondrial translational inhibition, antimycin A treatment and *anac017* overexpression, but not following ANAC017 loss of function in *rao2*.*1* (**Fig 3A, DataS6**). Increased ROS production due to limited electron transport chain capability could be the common cause of this retrograde signaling. Less assembled ETC Complex I was obvious in *lon1* while mitochondrial translation inhibition can limit mitochondrial-encoded subunits for efficient assembly. A relatively mild but significant UPR^cp^ was found specifically in *lon1* but not in other treatments or mutants (**Fig 3B, DataS6**). Notably, even a general down-regulation of UPR^cp^ target genes was observed for antimycin A treatment. Expression of several chloroplast chaperones were upregulated in *anac017* overexpression but also in *rao2*.*1*, suggesting their expression is induced by a noncanonical signaling pathway independent of ANAC017. It is still unclear what mediates this noncanonical signaling pathway to induce UPR^cp^. It is notable that Lon1 has been reported to be dual targeted to mitochondria and chloroplast (Daras et al., 2014). A specific UPR^cp^ induction suggests Lon1 might also function in chloroplast protein homeostasis alongside its known role in mitochondria. Compared to the apparent upregulation of UPR^mt^ and UPR^cp^, less than half the known set of UPR^er^ target genes were upregulated in *lon1* (**Fig 3C, DataS6**). Only ERF105, NAC103 and T6J4.21 showed consistent induction during mitochondrial stress and *anac017* overexpression while other known UPR^er^ target genes such as IRE1A were downregulated in the *lon1* line.

Ethylene has been proposed to be an important phytohormone for UPRs and mitochondrial retrograde signaling (Wang and Auwerx, 2017; He et al., 2022). Ethylene biosynthesis and signaling genes were found to be generally upregulated in *lon1* mutants. Ethylene overproduction and stimulated signaling appear to contribute to the short hypocotyl phenotype of *lon1* during skotomorphogenesis when optimal mitochondrial action is required (**Fig 4**). The short hypocotyl phenotype in *lon1* lines was partially recovered by ethylene signaling/biogenesis inhibitors AgNO_3_ and AVG. Ethylene can also inhibit cell elongation by an auxin-dependent and -independent pathway (**FigS8**). Cross-talk between ethylene and other phytohormones including auxin are potential regulators which cause small rosette, short root and growth retardation phenotypes in *lon1* in plants.

## Conclusion

Lon1 knockout inhibits mitochondrial-encoded protein translation due to formation of PPR and ribosomal protein aggregates. Protein aggregation can induce integrated cellular unfolded protein responses involving UPR^mt^, UPR^cp^ and partial UPR^er^. Collectively, this contributes to an integrated model to illustrate how plants stimulate integrated stress responses in response to mitochondrial translation rate changes to maintain balance in mitochondrial proteostasis. This evidence unites more features of Lon1 function across plants, yeast and animals and shows that the stunted plant phenotype of *lon1* is linked to hormone signaling rather than just energy metabolism.

## Methods

### Arabidopsis plants preparation

Arabidopsis plants were grown in soil under 16/8 hours light / dark cycle at room temperature (18-22 ° C), with a light intensity of 120-150 µ mol protons m^-2^ S^-1^. Arabidopsis mutants with a point mutation on Lon1 (*lon1-1*), a T-DNA insertion on Lon1 (*lon1-2*), and T-DNA insertion mutants *eto1-12* (AT3G51770, SALK_061581) and *ers2-1* (AT1G04310, SALK_203617) were grown under the same condition as *Col-0*. For hydroponically grown Arabidopsis, *Col-0* and *lon1* mutants seeds (30 mg) were surface sterilized in 70% ethanol (v/v) and then sterilized in 5% (v/v) bleach, 0.1% (v/v) Tween-20. Sterilized seeds were washed in sterile water five times before being dispensed into a plastic vessel containing 100 mL of growth medium (1/2 MS medium without vitamins, 1/2 Gamborg B5 vitamin solution, 5 mM MES, and 2.5% [w/v] Sucrose, pH 5.8). Plants were grown under long-day conditions on a shaker for 10-14 days. The continuous shaking allows Arabidopsis seedlings to be aerated allowing for respiration.

### Mitochondria isolation and protein aggregate enrichment

Mitochondria were isolated from 12-day-old hydroponically seedlings as described (Li et al., 2017b). A total of 30 μg of mitochondrial protein was dissolved in a lysis buffer (0.5% Triton X-100 (v/v); 30 mM Tris pH 7.4; 5 mM EDTA; 200 mM NaCl; 0.5 mM PMSF and protease Inhibitor cocktail). Mitochondrial lysates were centrifuged at 125,000 g for 45 min at 4°C to separate soluble protein supernatant and a pellet of aggregated proteins (Becker et al., 2012; Wilkening et al., 2018). Proteins in the supernatant were precipitated with nine times the volume of cold acetone. Protein pellets were resuspended in sample buffer and separated by SDS-PAGE (**FigS1**).

### Label-free quantification by mass spectrometry

30 μg mitochondrial protein isolated from *Col-0* and *lon1* mutant lines mitochondria were dissolve in resuspension buffer (7 M Urea; 2 M Thiourea; 50 mM NH_4_CO_3_; 10 mM DTT) and digested in-solution by trypsin. Protein aggregates were separated by 12% SDS-PAGE for 90 minutes using a high voltage at 180 V. Stained protein bands were excised and digested in-gel by trypsin. Tryptic peptides from in-gel/in-solution digestions were lyophilized in a Labconco centrifugal vacuum concentrator. C18 spin columns (Empore, 321069D) were used for desalting before LC-MS/MS analysis. Each sample was resuspended in 0.1% FA to a concentration of 1 µg/μl and made ready for mass spectrometry analysis. A Dionex Ultimate 3000 nano-HPLC system coupled with an Orbitrap Exploris 240 mass spectrometer (Thermo Fisher Scientific) were utilised for LC-MS/MS analysis. EASY-Spray columns (50 µm i.d. × 15 cm and 75 µm i.d. × 50 cm) home packed with 2 μm Thermo Scientific Acclaim PepMap RSLC C18 beads were used, with the following gradient profile delivered at 300 nL/min: 97% solvent A (0.1% formic acid in water) to 10% solvent B (0.08% formic acid in 80% acetonitrile) over 5 min, then 10% to 50% solvent B over 2 hours.

Raw data processing and database searches were performed with Proteome Discoverer software 2.4 (PD, Thermo Fisher Scientific). The Arabidopsis database TAIR10 from the Arabidopsis Information Resource (TAIR) was used to match peptide fragments resulting from the MS. PD Search parameters were: FDR<0.01 (strict) and FDR<0.05 (relaxed), carbamidomethyl (Cys) (static modification), Met oxidation, protein N-terminal modification acetylation (Arg, His, Ser, Thr, Tyr) and phosphorylation dynamic modification (Asp, Glu, His, Lys, Arg, Ser, Thr) with a maximum of 2 missed cleavage sites. Error tolerances of 10 ppm for MS and 0.02 Da for MS/MS.

### mRNA deep sequencing

Transcriptome sequencing and analysis were conducted by OE biotech Co., Ltd. (Shanghai, China) as a service. Total RNA of ten-day-old plate grown Arabidopsis seedlings under long day conditions was extracted using a RNA binding spin column. Purified total RNA were treated with DNase to remove contaminated genomic DNA. Oligo (dT) magnetic beads were utilized to enrich mRNA with poly (A) tails. RNA integrity was evaluated using the Agilent 2100 Bioanalyzer (Agilent Technologies, Santa Clara, CA, USA). The samples with RNA Integrity Number (RIN) ≥ 7 were subjected to the subsequent analysis. The libraries were constructed using TruSeq Stranded mRNA LT Sample Prep Kit (Illumina, San Diego, CA, USA) according to the manufacturer’s instructions. Then these libraries were sequenced on the Illumina sequencing platform (DNBSEQ-T7) and 125bp/150bp paired-end reads were generated. Raw data (raw reads) were processed using Trimmomatic (Bolger et al., 2014). The reads containing poly-N and low-quality reads were removed to obtain clean reads. Then the clean reads were mapped to the reference genome using hisat2 (Kim et al., 2015). The FPKM (Roberts et al., 2011) value of each gene was calculated using cufflinks (Trapnell et al., 2010), and the read counts of each gene were obtained by htseq-count (Anders et al., 2015). DEGs were identified using the DESeq (Anders and Huber, 2012) R package functions Estimate Size Factors and nbinomTest P-value < 0.05 with fold change >1.5 was set as the threshold for significantly differential expression. Hierarchical cluster analysis of DEGs was performed to explore the gene expression patterns. GO enrichment and KEGG (Kanehisa et al., 2008) pathway enrichment analysis of DEGs were respectively performed using R based on the hypergeometric distribution. RNA-seq data are summarized in **DataS5** and can be accessed with GEO number GSE216535.

### Analysis of in organelle protein synthesis

An in-organelle protein synthesis assay was performed following a procedure as described (Kwasniak-Owczarek et al., 2022). A translation mix (5 mM KH_2_PO_4_, 2 mM guanosine 5’-triphosphate, 0.4 M mannitol, 60 mM KCl, 2 mM DTT, 50 mM HEPES, 10 mM MgCl_2_, pH 7.0) was divided into two equal volumes for mitochondrial translation and bacterial control. 25 μL amino acid mixture minus L-methionine (from 1 mM stock) and 1 mg BSA were supplied to 1 ml translation mix. Freshly isolated mitochondria (150 µg proteins) were added to 100 µl of mitochondrial translation mix (supplemented with 10 mM malic acid, 1 mM pyruvate, 4 mM adenosine-5-diphosphate and bacterial control translation mix (supplemented with 20 mM sodium acetate). 30 µCi [^35^S]-Met were supplied to each sample and they were incubated at 25 °C for 60 min with rotation. The reaction was stopped by supplying 350 µl of ice cold 1x Wash Buffer (0.6 M sucrose, 20 mM TES, pH 7.5) containing 10 mM L-methionine and50 µg/mL puromycin. Mitochondrial proteins were separated by SDS-PAGE. Stained SDS-PAGE bands were excised and transferred to scintillation vials and emulsified in 10 ml of Ultima Gold XR scintillation liquid (Perkin Elmer) in darkness at room temperature overnight. Counts Per Minute (CPM) of the [^35^S] radioactivity was measured using a liquid scintillation counter (Tri-Carb 4810 TR; PerkinElmer).

### RNA extraction and Q-PCR analysis

Seedlings of ten-day-old plant grown seedlings of *Col-0* and two *lon1* mutant lines grown under long day conditions were collected for RNA extraction. Collected seedlings (∼0.1g) were snap frozen in liquid nitrogen and homogenized to powder using beads (2 mm) by a homogenizer. RNA was extracted using TaKaRa MiniBEST Plant RNA Extraction Kit (TaKaRa, 9769) following the manufacturer’s instructions. 500ng of RNA was used for cDNA synthesis with TSINGKE Goldenstar™ RT6 cDNA Synthesis Kit Ver.2 (TSINGKE, TSK302M). Transcripts of selected genes were quantified using TB Green® *Premix Ex Taq*™ (Tli RNaseH Plus) (TaKaRa, RR420A) with Realplex2 (Eppendorf). Q-PCR data were normalized to a housekeeping gene *UBQ10* before being compared across genotypes.

### Ethylene emission measurements

Etiolated Arabidopsis seedlings, one to five days after sowing were incubated in sealed vials with 3-ml 1/2 MS (Murashige and Skoog) liquid medium and placed in a plant growth chamber for 24 hours (18-22 °C) in darkness. Ethylene emission in the vials were measured by gas chromatography (Agilent 7890A) as described (Xu et al., 2021).

## Supporting information

DataS1

DataS2

DataS3

DataS4

DataS5

DataS6

DataS7

## Open accessible data

PRIDE Project Name: Arabidopsis Lon1 disruption inhibits mitochondrial translation and induces retrograde signals to rebalance mitochondrial protein homeostasis (total mitochondria)

Project accession: PXD038026

PRIDE Project Name: Arabidopsis Lon1 disruption inhibits mitochondrial translation and induces retrograde signals to rebalance mitochondrial protein homeostasis (protein aggregations)

Project accession: PXD038029

RNA-seq data can be accessed through the following link https://www.ncbi.nlm.nih.gov/geo/query/acc.cgi?acc=GSE216535

## Acknowledgements

Dr. Oliver Berkowitz and Professor Jim Whelan are thanked for advice on RNA seq processing. Dr. Congcong Lu and Dr. Yuzhen Wu in Nankai Proteomics Centre are thanked for assistance for mass spectrometry data collection and analysis. Professor Jing Zhang and Ms. Xiaoyu Wang are thanked for assistance in Arabidopsis hypocotyl staining.

## Author Contribution

LL, NNW and AHM designed the research; CS performed plant culture and biochemical experiments; CS performed ethylene measurement together with CX. Proteomics mass spectrometry and analysis was performed by YYL, YQH and CS. MMY, TTL and YYL prepared plant materials and performed mitochondria preparation. CS and JJL performed in organelle protein synthesis assay. LL, CS and AHM contributing to the writing and revision of the article.

## Supplemental Figures

**FigS1.**
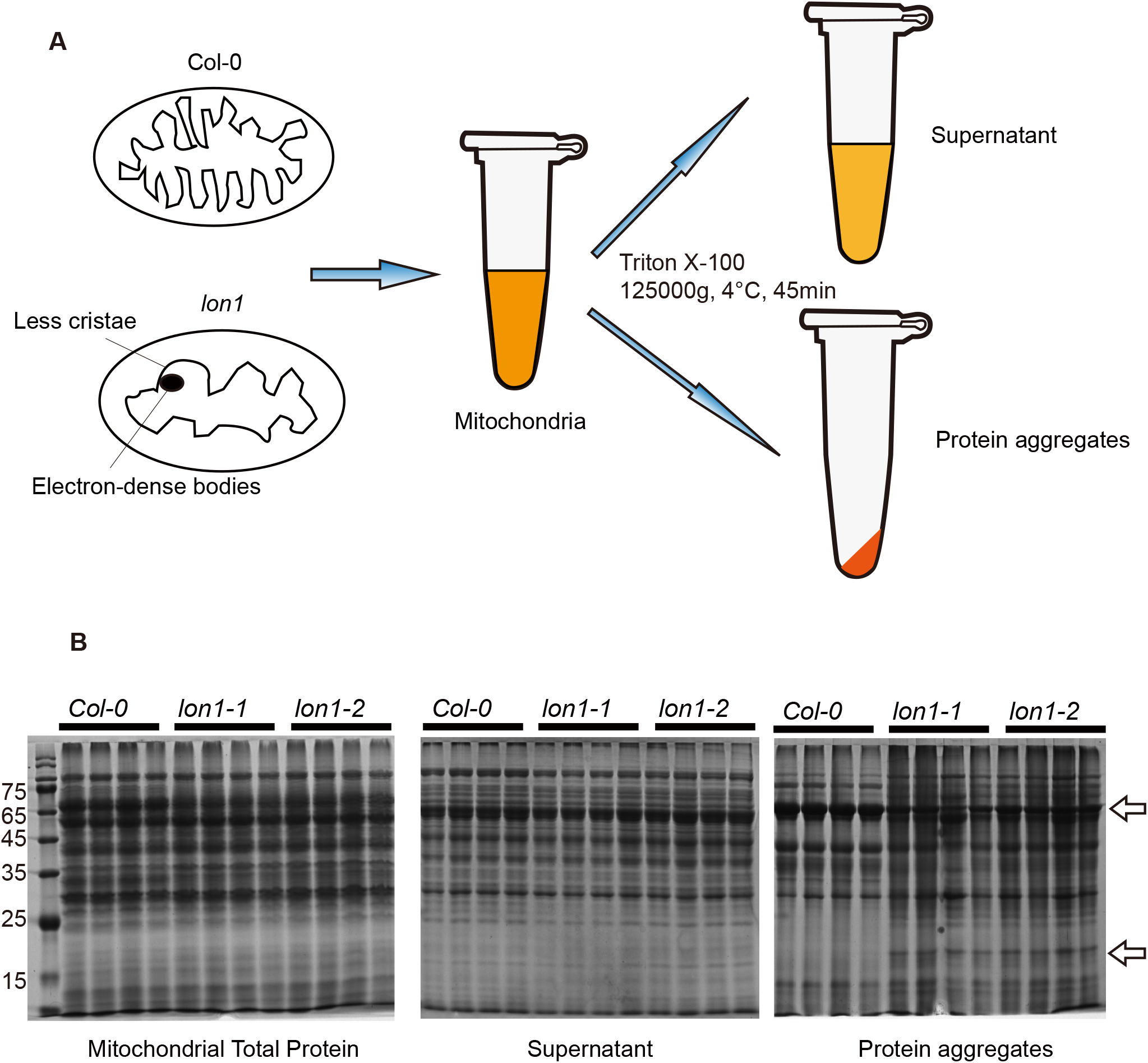
Mitochondrial protein aggregates separation by difference in detergent solubility. The workflow to fractionate protein aggregates in *Col-0* and *lon1* mutant lines **(A)**. 30 µg supernatant/aggregated proteins were separated by SDS-PAGE and stained by Coomassie blue for visualisation **(B)**. Arrows show representative bands with different patterns in *Col-0* and *lon1* mutant lines. Supports Fig 2.

**FigS2.**
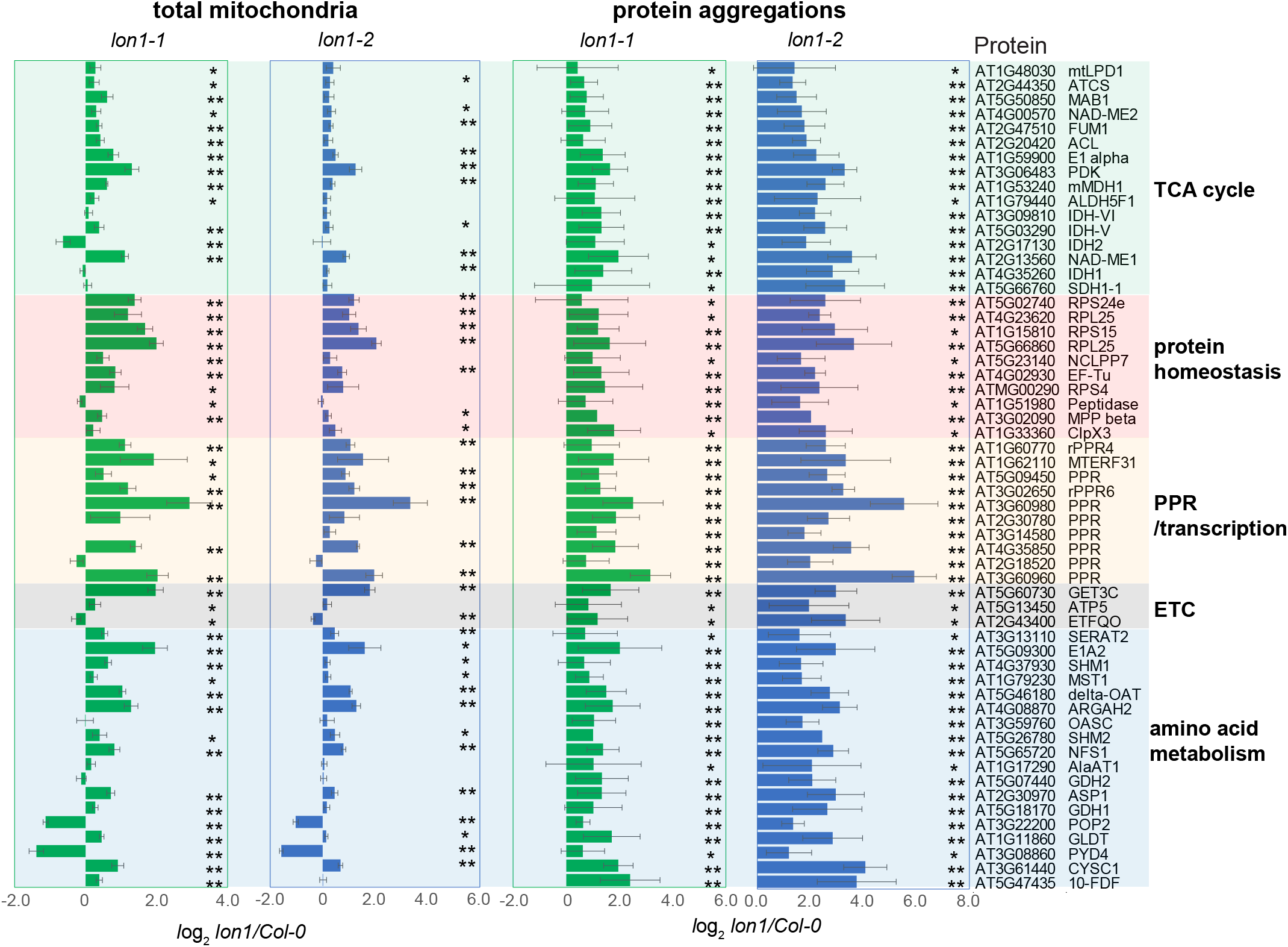
Protein abundance changes in total mitochondria and protein aggregates. Fifty-seven proteins with significant accumulation in at least one of the *lon1* mutant lines and an average aggregation factor over 1.5 fold are shown. A Student’s T-test was utilised for statistical analysis (*P<0.05, ** P<0.01). Supports Fig 2.

**FigS3.**
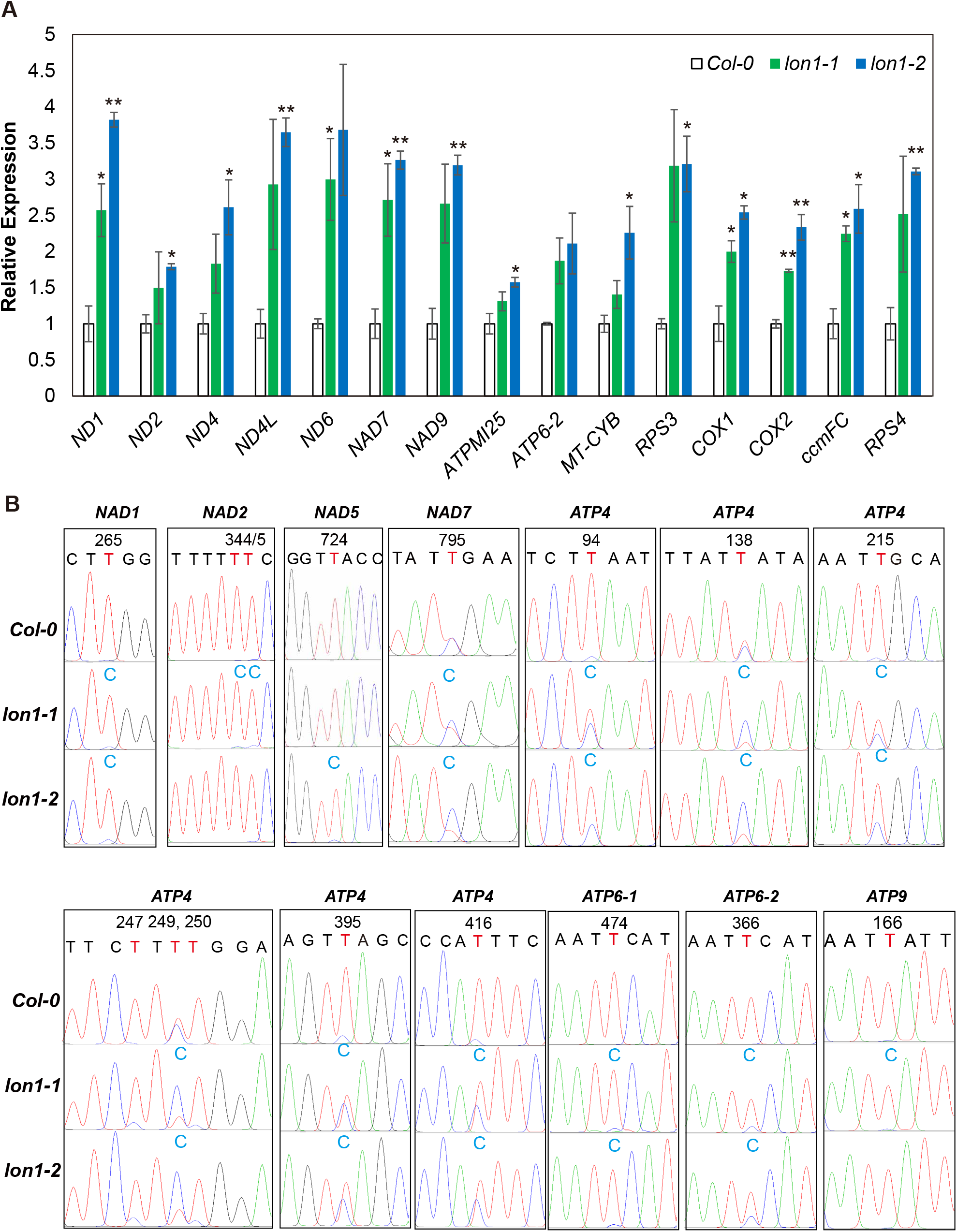
Mitochondrial-encoded gene expression and RNA editing. Transcript levels of fifteen mitochondrial-encoded genes measured by qRT-PCR using *UBQ10* as an internal control (**A**). A Student’s T-test was utilised for statistical analysis (*P<0.05, ** P<0.01). Nine genes with known RNA editing sites were sequenced. Sequencing chromatograms of eight genes with observable overlapped T/C peaks are provided (**B**). All results are supplied in DataS4. Supports Fig 2.

**FigS4.**
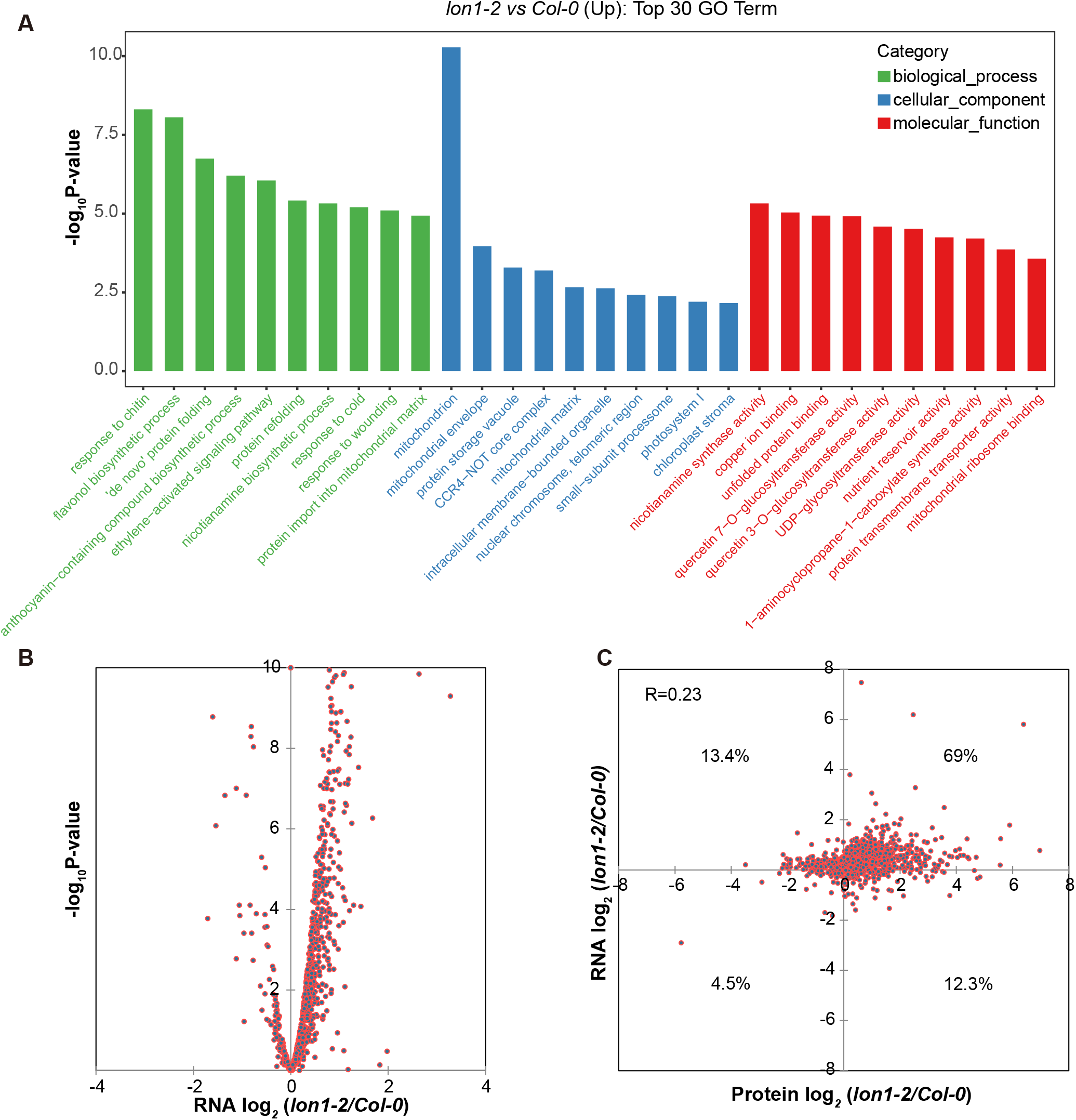
Lon1 disruption induces gene expression of chaperones and mitochondrial localised proteins. GO enrichment analysis was performed using 4662 upregulated genes (FC>2, P<0.05) as target and all encoding genes as background. The top 30 enriched GO terms for biological process, cellular components and molecular function were shown (**A**). Log-transformed fold change for 921 genes which were also identified in protein analysis in both *lon1-2* and *Col-0* were visualised by volcano plotting (**B**). Log-transformed transcript level of 921 genes and their protein abundance were visualised by dot plotting (**C**). Supports Fig 3.

**FigS5.**
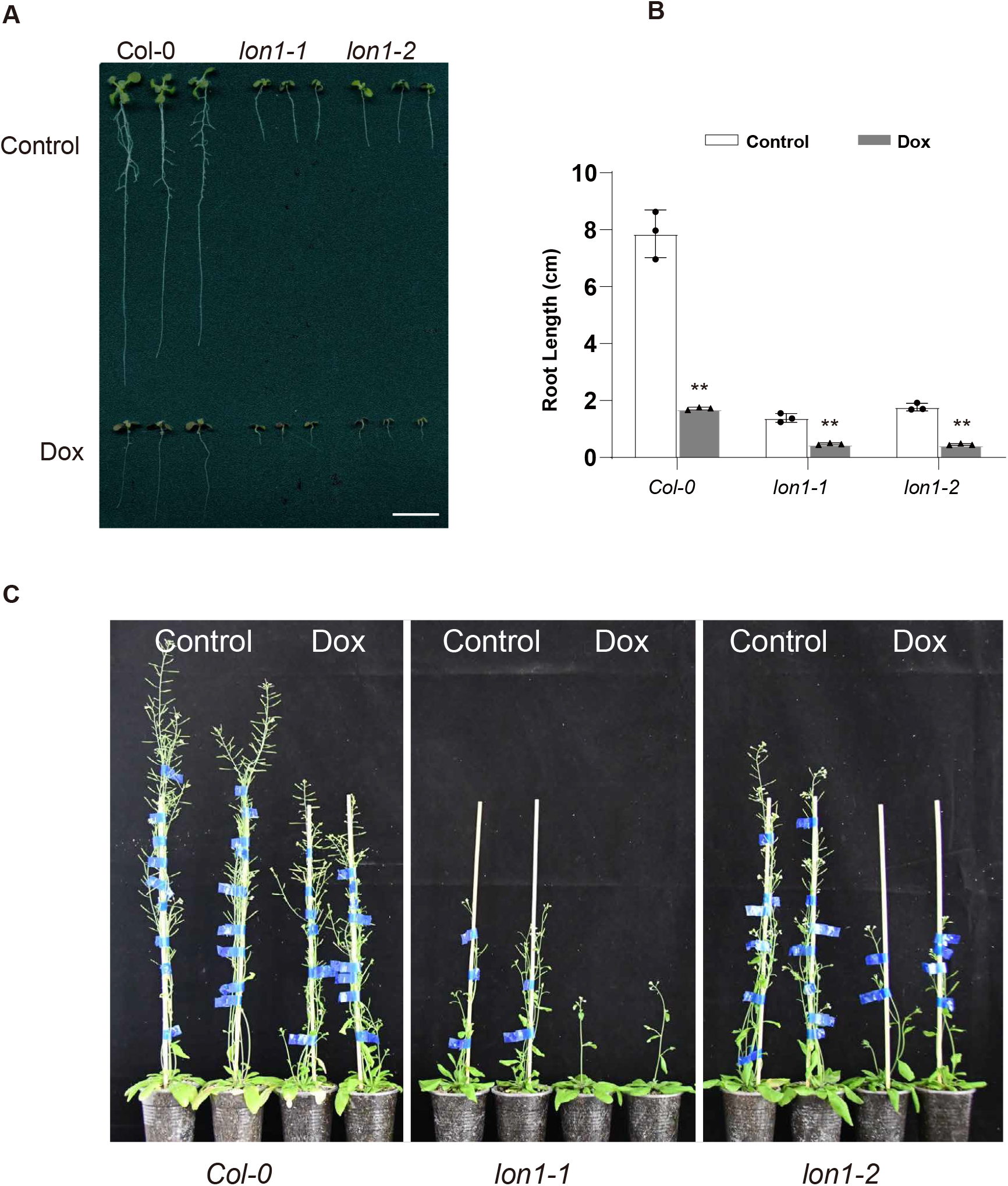
Doxycycline treatment induces Arabidopsis short root, small rosette and growth retardation in *Col-0* and *lon1* mutants. Two-week-old seedlings grown on MS plates with/without doxycycline (Dox) (**A**). Root length was measured and shown in column graphs (**B**). The ruler bar is 1 cm. Error bars show standard deviation of three biological replicates. A Student’s T-test was utilised for statistical analysis (*P<0.05, ** P<0.01). Twenty-four-day-old Arabidopsis plants grown in soil and watered with water supplied with/without 25 mg/mL Dox (**C**). Supports Fig 3.

**FigS6.**
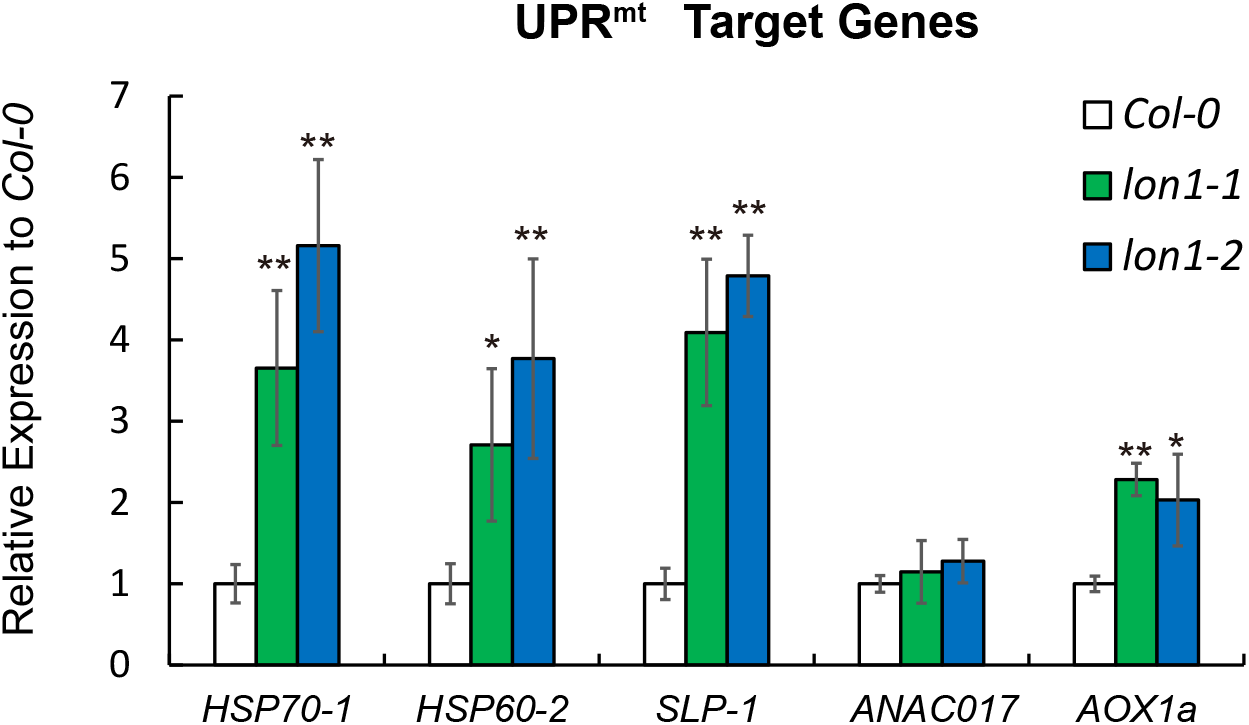
Validation of UPR^mt^ gene expression by Q-PCR. Abundance of transcripts of selected UPR^mt^ target genes (*HSP70-1, HSP60-2, SLP-1, ANAC017, AOX1a*) were measured by quantitative real-time PCR. Error bars show standard deviations of three biological replicates. A Student’s T-test was utilised for statistical significance (*P<0.05, **P<0.01). Supports Fig 3.

**FigS7.**
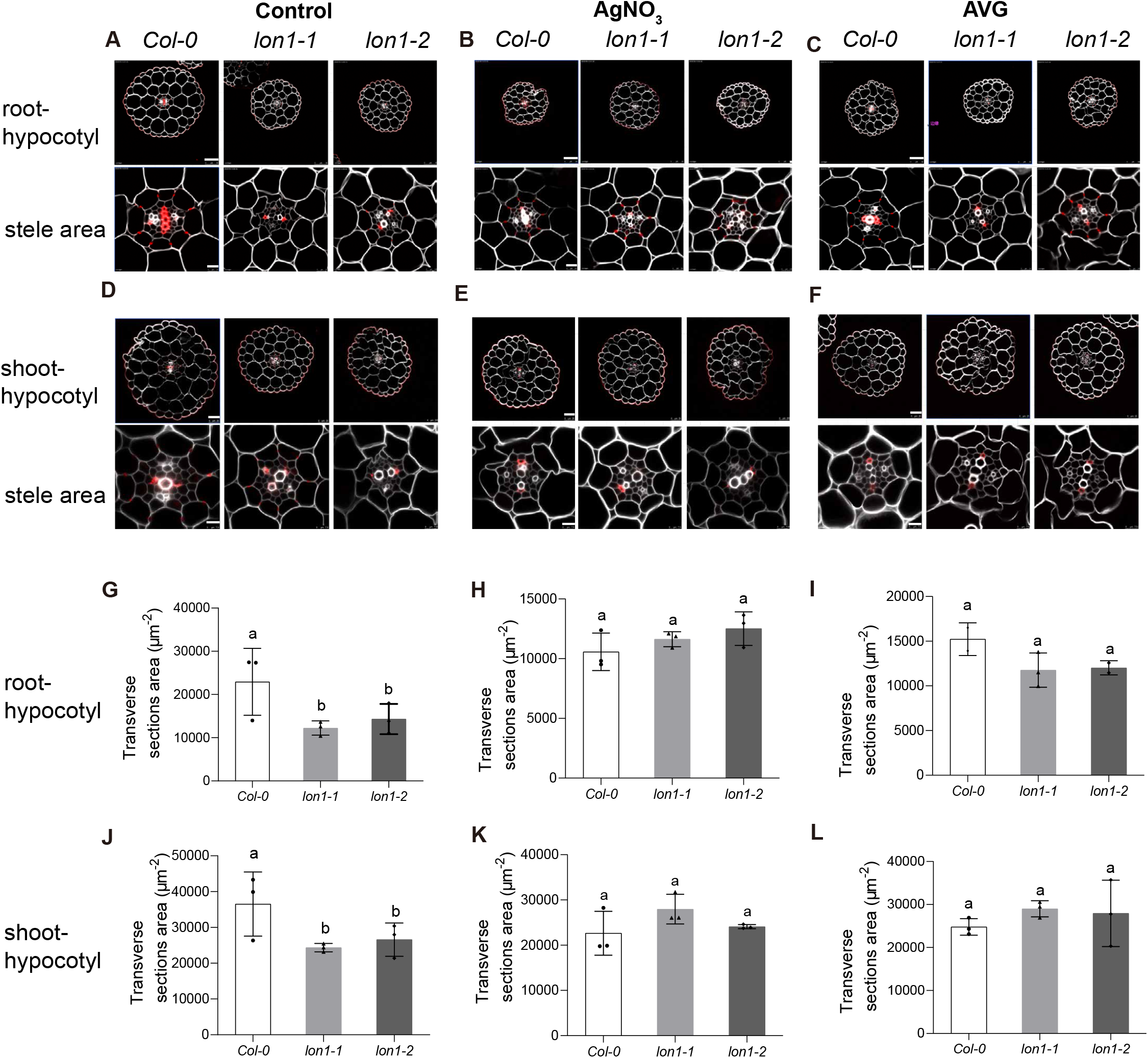
Lon1 defects cause slimmer hypocotyl and retarded lignin biogenesis. *Cell*ulose and lignin in five-day-old Arabidopsis hypocotyl transverse section (control, AgNO_3_ and AVG treatments) revealed by SCRI Renaissance (cellulose-white) and Basic Fuchsin (lignin-red) staining. Root-hypocotyl and shoot-hypocotyl are shown separately (control-**A, D**; AgNO_3_-**B, E** and AVG-**C, F)**. Casparian strips and xylem vessels rich in lignin are red after staining. Transverse section areas are shown in column graphs (control-**G, J**; AgNO_3_-**H, K** and AVG-**I, L**). Error bars show standard deviations of measurements of three etiolated Arabidopsis plants. The ruler bar is 50 μm for transverse sections and 10 μm for zoomed in stele areas. Supports Fig 4.

**FigS8.**
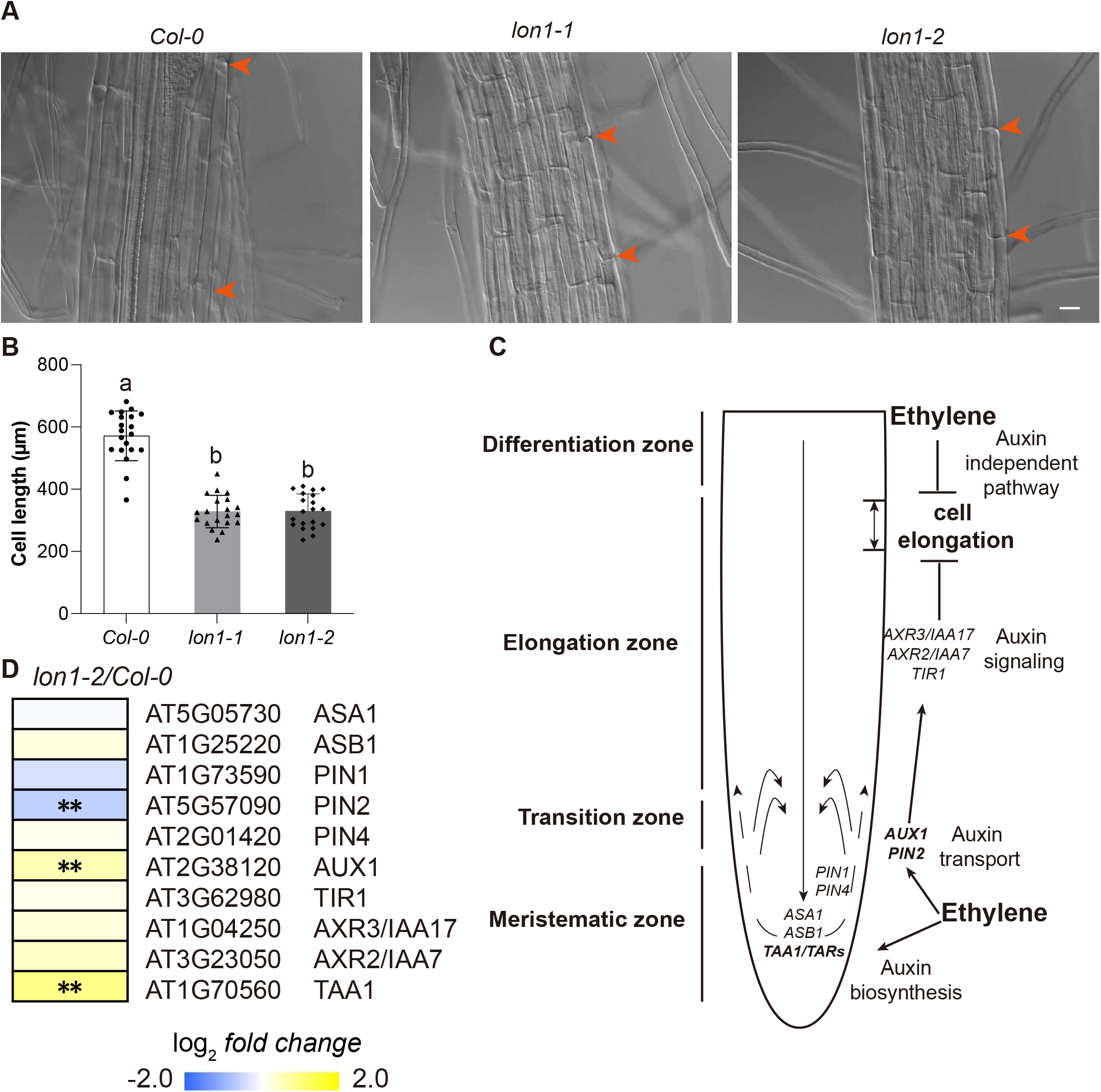
Ethylene overproduction in *lon1* mutants affect root cell elongation through auxin dependent/independent pathways. Differentiation zone root cells of seven-day-old *Col-0, lon1-1* and *lon1-2* Arabidopsis were visualised by microscopy (400x, Ruler=20 μm). *Cell* length was measured and compared between *Col-0* and *lon1* mutants (**A**). Orange arrowheads indicate cell boundaries. Error bars show standard deviations (n=20). Grouping of cell length was determined by one-way ANOVA analysis (**B**). Auxin-associated gene expression (according to a log2 fold change cut-off) in *lon1-2* compared to *Col-0* are shown in heatmaps (**C**). A Student’s T-test were used for statistical analysis (*P<0.05, **P<0.01). Ethylene inhibits root elongation through auxin dependent/independent pathways (Hu et al., 2017). TAA1, AUX1 and PIN2 involved in auxin biosynthesis and transport with significant changes were highlighted using bolded fonts in the model (**D**).

## Supplemental Data

DataS1 Mitochondrial protein abundance measurement by label free mass spectrometry.

DataS2 Protein aggregate protein abundance measurement by label free mass spectrometry.

DataS3 Seventy-eight proteins with average aggregation factor over 1.5 fold.

DataS4 Editing efficiency of seventy sites in nine mitochondrial-encoded genes.

DataS5 RNA deep sequencing of *lon1-2* and *Col-0*.

DataS6 UPR genes in *lon1* and reported mitochondrial retrograde signaling associated treatments or mutant lines.

DataS7 Combined analysis of transcripts and protein abundance of mitochondrial localised proteins.

